# The role of non-additive gene action on gene expression variation in plant domestication

**DOI:** 10.1101/2022.10.13.511672

**Authors:** Erik Díaz-Valenzuela, Daniel Hernández-Ríos, Angélica Cibrián-Jaramillo

## Abstract

**Background:** Plant domestication is a remarkable example of rapid phenotypic transformation of polygenic traits such as organ size. Evidence from a handful of study cases suggests this transformation is due to gene regulatory changes that result in non-additive phenotypes. Employing data from published genetic crosses, we estimated the role of non-additive gene action in the modulation of transcriptional landscapes in three domesticated plants: maize, sunflower, and chili pepper. Using A. *thaliana*, we assessed the correlation between gene regulatory network (GRN) connectivity properties, transcript abundance variation, and gene action. Finally, we investigated the propagation of non-additive gene action in GRNs.

**Results:** We compared crosses between domesticated plants and their wild relatives to a set of control crosses that included a pair of subspecies evolving under natural selection and a set of inbred lines evolving under domestication. We found abundance differences on a higher portion of transcripts in crosses between domesticated-wild plants relative to the control crosses. These transcripts showed non-additive gene action more often in crosses of domesticated-wild plants than in our control crosses. This pattern was strong for genes associated with cell cycle and cell fate determination, which control organ size. We found weak but significant negative correlations between the number of targets of trans-acting genes (Out-degree) and both the magnitude of transcript abundance differences a well as the absolute degree of dominance. Likewise, we found that the number of regulators that control a gene’s expression (In-degree) is weakly but negatively correlated with the magnitude of transcript abundance differences. We observed that dominant-recessive gene action is highly propagable through GRNs. Finally, we found that transgressive gene action is driven by trans-acting regulators showing additive gene action.

**Conclusions:** Our study highlights the role of non-additive gene action on modulating domestication-related traits such as organ size via regulatory divergence. We propose that GRNs are shaped by regulatory changes at genes with modest connectivity, which reduces the effects of antagonistic pleiotropy. Finally, we provide empirical evidence of the propagation of non-additive gene action in GRNs, which suggests a transcriptional epistatic model for the control of polygenic traits such as organ size.

## Background

Plant domestication has been widely employed as a model to study phenotypic evolution [1]. One of the most remarkable morphological features that distinguish a domesticated from its wild progenitor is the gigantism of vegetative and reproductive organs [1–3]. Studies employing genetic crosses between domesticated and wild plants have revealed this gigantism is a recessive feature, as the domesticated phenotype is often masked in F1 hybrids [4–6]. This evidence suggests that genetic factors underlying domestication-related phenotypes might include loss-of-function (LOF) mutations whose function can be complemented by a wild functional allele. Furthermore, it has been reported that these domestication-related phenotypes are often disadvantageous or even lethal in natural populations, suggesting these recessive phenotypes are likely the product deleterious mutations [7]. Empirical work revealed these loss-of-function mutations often affect the regulation of genes rather than their coding products [2, 6, 8]. Which suggests a relevant role of transcript abundance variation in the modulation of domestication-related phenotypes. Indeed, several studies have reported that variation on transcript abundance of genes involved in organ size tends to show a recessive pattern of gene action for the domesticated phenotype [9, 10]. Transcript abundance is controlled by regulatory interactions between cis-acting non-coding DNA sequences and trans-acting factors such as transcription factors and chromatin remodeling factors [11]. These interactions between regulatory elements act coordinately within the cells forming genetic circuits or gene regulatory networks (GRNs) to determine quantitative phenotypes such as organ size [12].

Genes coding for cell cycle and cell fate determination are among relevant members of GRNs controlling meristem and therefore organ size, as they control the rates of cell division and cell expansion [13]. It is likely that mutations affecting the regulatory activity trans-acting factors with a larger number of transcriptional targets produce greater pleiotropic effects than genes connected with a few genes [14].

Hence, it is possible that the former experience greater selective constraint to avoid antagonistic pleiotropy (negative effects on fitness due to regulatory variation on multiple target genes). Similarly, genes that are regulated by a large number of trans-acting factors would experience selective constraint as they have key roles in development [15].

In diploid organisms, cis-acting mutations often produce additive gene action. A mechanistic explanation for this pattern is that cis-acting mutations act in an allele-specific fashion, so the expression of one allele does not affect the expression of the other and the resulting phenotype is the average of the parental alleles. Conversely, trans-acting factors bind to cis-regulatory elements of the two alleles, modulating the phenotype towards one parent, and thus producing non-additive (dominant-recessive) gene action [16, 17]. As mentioned above, recessive phenotypes are often associated with loss-of-function mutations. It is therefore possible that trans-acting factors highly connected in GRNs would display dominant recessive gene action less often to avoid antagonistic pleiotropy. It is also possible thus, that trans-acting factors showing dominant-recessive gene action would produce dominant-recessive gene action on their targets downstream. Some simulation-based studies have revealed that dominant-recessive gene action within GRNs is highly propagable [18]. The role of non-additive gene action on transcript abundance differences between wild and domesticated plants has been assessed in independent studies [9, 19–24]. However, the use of different analytical approaches to both quantify transcript abundance and measure gene action has yielded contrasting results that are difficult to compare among them. Furthermore, besides a few studies on *Drosophila* and yeast that have assessed the association between GRN connectivity properties and transcript abundance differences, no studies have approached such associations in plant models.

In this study, we evaluated the contribution of gene action to transcript abundance divergence in plants evolving under domestication using species that exhibit gigantism of reproductive organs. These species include maize (*Zea mays*), chili pepper (*Capsicum annuum*) and sunflower (*Helianthus annuus*). Although they exhibit different morphology, human-mediated selection has resulted in the enlargement of their reproductive organs such as flowers, fruits and seeds [2]. Because they were domesticated in what it is nowadays Mexico and the United States, they were likely exposed to a common set of cultural preferences and agricultural practices [1]. As control scenarios of selective pressures, we included data from crosses of two switchgrass (*Panicum hallii*) subspecies evolving under natural selection as well as two inbred lines of rice (*Oryza sativa*) and thale cress (*Arabidopsis thaliana*).

We hypothesized additive gene action on transcript abundance to be less frequent in the genetic crosses evolving under domestication than in the crosses evolving under natural selection. We particularly predicted this scenario for genes associated with cell cycle and cell fate determination because they are directly associated with the gigantism of reproductive organs. Using a co-expression based GRN, we also investigated the association between both the effect size of transcript abundance differences and gene action to GRN connectivity properties (In-degree and Out-degree) in *Arabidopsis thaliana* F1 hybrids, to test for evidence of evolutionary constraint in the expression of genes with high Out-degree and high In-degree. Additionally, to assess the role of transcriptional epistasis in propagating gene expression variability in GRNs, we analyzed the similarity of gene action between trans-acting regulators and their target genes. We observed an overall reduction of additive gene action on the domesticated transcript abundance phenotype, with a marked pattern for transcripts associated with cell cycle and cell fate determination. We also observed a negative correlation between GRN connectivity properties and both effect size of transcript abundance differences and gene action. Finally, we observed that dominant-recessive gene action is highly propagable through GRNs and that trans-acting factors showing additive gene action might be driving transgressive gene action on genes downstream.

## Results

### The genome-wide patterns of inheritance of transcript accumulation unravel an overall reduction of additive gene action in domesticated phenotypes

To investigate the genetic nature of phenotypic variation at the transcript abundance level in the context of plant domestication, we compiled a large data set that included genetic crosses evolving under natural selection and under domestication (**Table S1**). Our data set includes the genome-wide transcriptomes of both F0s and F1 hybrids of two switchgrass (*Panicum hallii*) subspecies diverging under natural selection; of cultivated maize (*Zea mays*), chili pepper (*Capsicum annuum*), and sunflower (*Helianthus annuus*) genotypes as well as their wild progenitors; of two rice (*Oryza sativa*) inbred lines and two reciprocal crosses of thale cress (*Arabidopsis thaliana*). We were particularly interested in transcripts showing abundance differences between the F0s of each genetic cross because as we see it, they are a proxy of organismal phenotypic variation. We observed a wide range (1% to 44%) in the portion of the transcriptome showing significant abundance differences between F0s (p-adjusted < 0.05, |Log2(FC)| > 0.5) (**Table S1**). This difference was substantially larger in the comparisons between domesticated-wild F0s (average of 35%) compared to the switchgrass F0s (1%), which diverged under natural selection. This result could be partially explained to the nature of the grass itself, or by differences in sequencing coverage. However, a random sampling-based coverage analysis showed a uniform median transcript coverage of ~60X for all the libraries used in this study (**Fig S1**), discarding this feature as a possible source of bias in our results. Therefore, this result is instead consistent with the polygenic nature of domestication-related phenotypes such as the overall increase in organ size, which requires the coordinated expression of large constellations of developmentally-regulated genes [2, 25]. We employed the degree of dominance (*k*) as a proxy of gene action to investigate the nature of genetic variation controlling transcript abundance variation across the twelve genetic crosses. The degree of dominance (*k*) quantitatively describes the extent to which the F1 hybrid phenotype deviates from a midpoint between the two F0 progenitors. Usually, *k* values near zero indicate purely additive gene action or equal contributions of the two alleles to the F1 hybrid phenotype. *k* values around |1| reflect dominant-recessive gene action, in which the phenotype produced by one allele masks the phenotypic expression of the other allele. While *k* values above |1.25| are interpreted as transgressive gene action. In this mode of gene action, the F1 hybrid displays a phenotype above (overdominance) or below (underdominance) the phenotypic values of its F0 progenitors. We hypothesized additive gene action on transcript abundance to be less frequent in the genetic crosses evolving under domestication than in the switchgrass genetic cross. Overall, our results show that the vast majority of the transcripts show *k* values within −5 and 5 (**Fig 1A**), this range is consistent with *k* estimates on plant morphology and transcriptional phenotypes in maize and chili pepper, respectively [9, 26]. Supporting our hypothesis, the data of switchgrass show a symmetrical shape with a mode centered near to zero, within the range of additive gene action (|*k*| < 0.25). Whereas for the crosses of domesticated-wild plants and inbred-inbred genotypes, the distributions show non-symmetrical shapes with modes outside the additive gene action range but below the transgressive gene action range (|*k*| > 0.25, |*k*| < 1.25), suggesting a relevant role of partially and fully dominant-recessive allelic interactions in shaping transcript abundance under domestication (**Fig 1A**).

**Figure 1.**
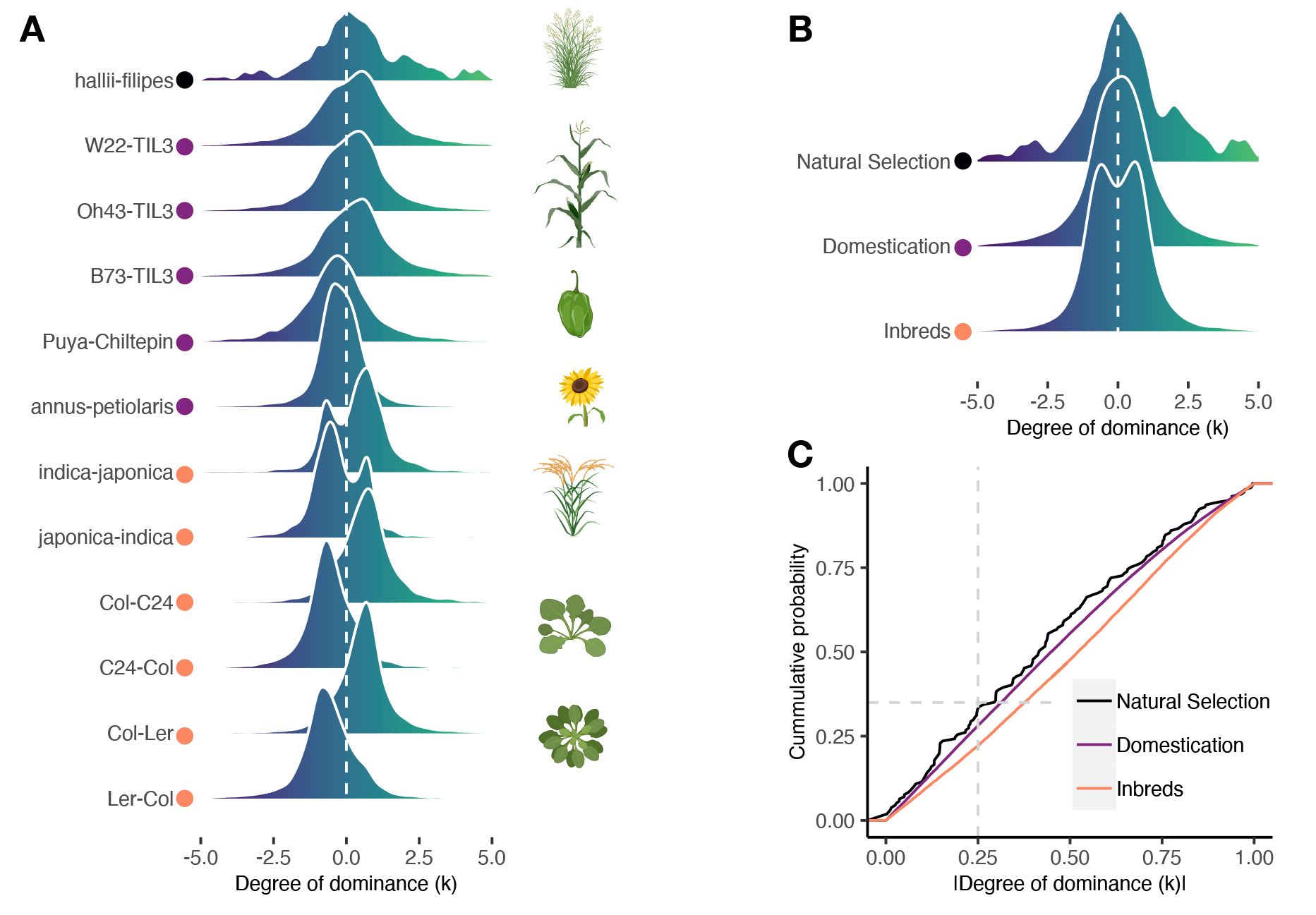
Inheritance of transcript abundance unveils a relevant role of non-additive gene action in domesticated plants. **A**) Distribution of the degree of dominance (*k*) across a series of genetic crosses including switchgrass, maize, chili pepper, sunflower, rice and Arabidopsis. |*k*| > 0.25 indicate non-additive inheritance. The color of circles indicates the category of the genetic cross. **B**) Distribution of *k* for each category of genetic cross. Natural Selection: wild-wild; Domestication: cultivated-wild; Inbreds: cultivated-cultivated. **C**) Empirical cumulative distribution analysis revealed the switchgrass species evolving under natural selection are distinguishable from those evolving directional selection for both Domestication (Kolmogorov-Smirnov: D = 0.21; p-value = 2.8E-14) and Inbreds (Kolmogorov-Smirnov: D = 0.28; p-value = 2.2E-16) genetic crosses.

In particular, we noticed that in the domesticated-wild crosses of chili pepper and sunflower, the domesticated phenotype tended to be recessive, although the opposite pattern was shown in maize-teosinte crosses [27]. Finally, transgressive gene action (|*k*| > 1.25) was a common feature for all the genetic crosses.

Data for *k* were aggregated into three discrete categories of genetic crosses, namely: natural selection, domestication, and inbreds, and their distributions were compared using a probability density function (**Fig 1B**). In general, we observe a pattern of wider distributions in domestication and inbreds compared to natural selection, supporting our prediction of less of additive gene action under domestication.

To formally test this observation, the distributions were compared using a cumulative distribution approach (**Fig 1C**). We observed a significantly larger portion of transcripts showing additive gene action (35% of total) in the cross of natural selection compared to the domestication and inbred crosses (25% and 20%; Fisher’s exact test: p = 2.905e-5, p = 5.996e-11, respectively).

By employing a pairwise Kolmogorov-Smirnov test to quantitatively compare the distributions, a substantial and significant difference was found for the three comparisons (Natural selection vs Domestication: p-value = 2.864e-14; Natural selection vs Inbreds: p-value = 2.2e-16; Domestication vs Inbreds: p-value = 2.2e-16). A similar result of less additive gene action in domesticated plants was observed by comparing gene action between cultivated maize and wild teosinte populations [28].

### Cell cycle and cell fate determination-related genes tend to display non-additive gene action in genetic crosses involving domesticated plants

A gene ontology (GO) enrichment analysis was performed across the twelve genetic crosses to gain insight into the biological functions associated with transcriptional differences and mode of gene action. Instead of testing for the enrichment of GO terms in the sets of differentially abundant transcripts in each genetic cross, we tested whether a set of specific GOs, likely related to the domestication syndrome were enriched or present in the discrete categories of gene action. Overall, we observed that the majority of the GOs are represented by sets of transcripts showing non-additive gene action This supports the role of non-additive gene action in producing phenotypic divergence at the transcript abundance level, particularly in domesticated-wild genetic crosses. Among the most relevant ontologies showing this pattern, we found cell cycle (GO:0007049), cell fate determination (GO:0001709), flower development (GO:0009909), response to auxin (GO:0009733), and seed dormancy (GO:0009793), which could be related to the domestication syndrome (**Fig S2**). We found weak enrichment for some of those categories, but this result was expected because the number of genes in each category is a subsample of the total DE transcripts and hypergeometric tests are sensitive to small sample sizes [29]. We consider the presence-absence variation of GOs in each category of gene action provides relevant evidence of which biological functions were altered in each divergence type. These results highlight the role of non-additive gene action in modulating cellular and development-related phenotypes through transcriptional divergence under domestication. The genetic cross of switchgrass, evolving under natural selection, shows abundance differences only on transcripts associated with nitrogen metabolism (GO:0006807), which reflects local adaptation to the environmental variation between the two studied ecotypes [29]. The continuous distributions of the degree of dominance (*k*) were analyzed for genes associated with meristematic activity such as the cell cycle, cell fate determination, flower development, auxin response, and seed dormancy, which are strongly related to the domestication syndrome. In the switchgrass cross, in addition to have found a small number of genes with differential expression performing these biological functions, these genes show values of *k* closer to zero, indicating additive gene action (**Fig 2**).

**Figure 2.**
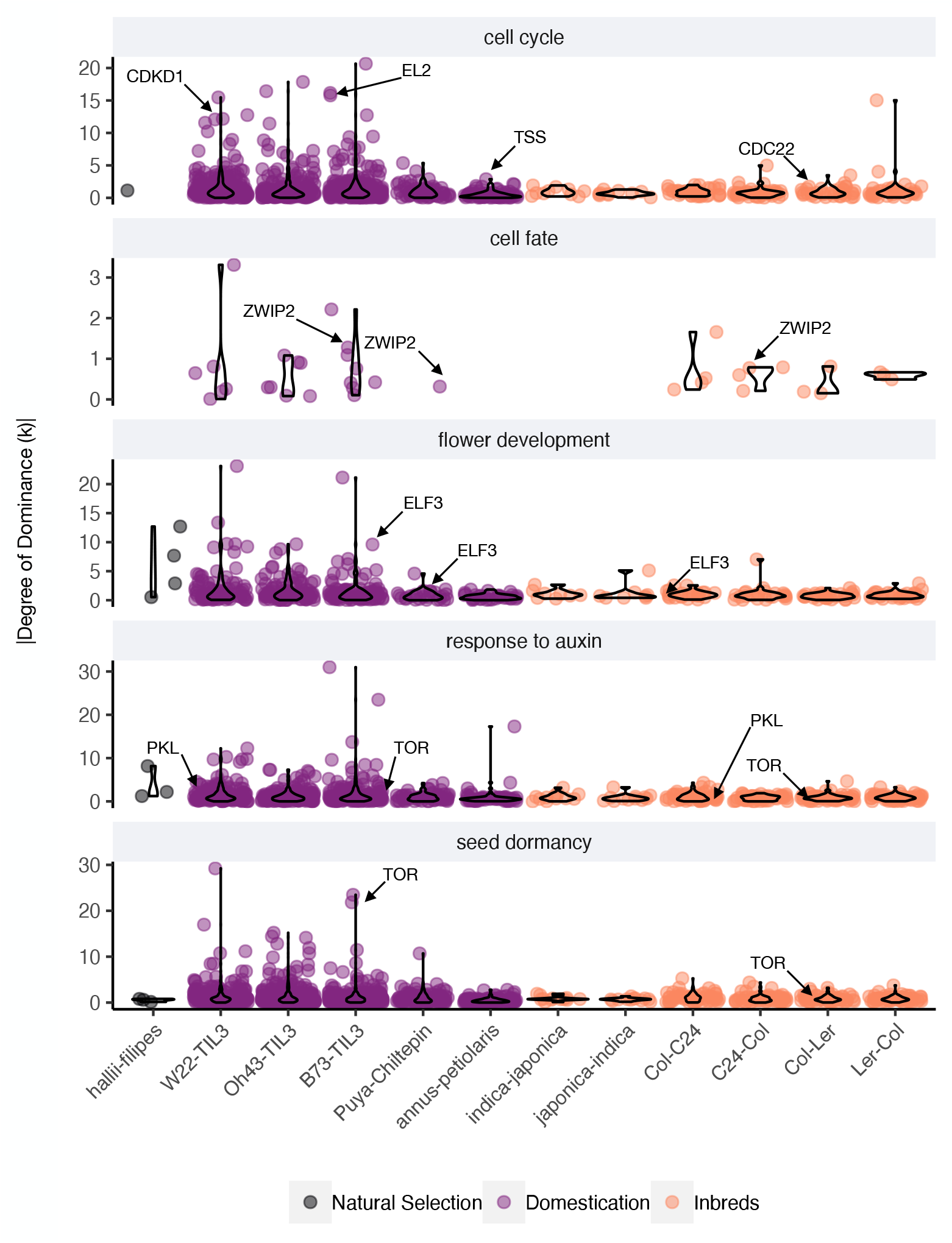
Inheritance of meristem-related genes in cultivated plants suggest selection under domestication employs non-additive gene action. Jitter + violin plots show the distribution of the absolute degree of dominance (*k*) for genes annotated to each biological process. Labels denote specific genes with experimental evidence of their roles in each biological process.

Congruently, for the genes associated with the domestication syndrome, we observed that crosses involving domesticated and inbred plants showed higher *k* values compared to the cross evolving under natural selection. Further, we found that the median *k* values for transcripts annotated with those biological functions showed gene action that is distinguishable from additive gene action (|k > 0.25|) (**Fig 2, Table S2**). This result suggests that genes associated with meristematic activities tend to display dominant-recessive or transgressive gene action. *CDKD1*, *CDC22, EL2*, and *TSS* were amongst the most relevant genes annotated as cell cycle regulators. They are part of the machinery that ensures the transitions between different cell cycle stages. For example, *CDKD1* encodes for a cyclin-dependent kinase that controls the transition between the G2 and M phases of the cell cycle [30]. CDC22, a cofactor of the anaphase-promoting complex (APC) controls meristem size by regulating mitosis [31]. While EL2 inhibits the progression of the cell cycle by directly binding to CDKD1, thus having a role in the endoreduplication cell cycle and cell size [32]. Finally, TSS indirectly controls the cell cycle and organ size by modulating sugar availability, therefore having a relevant role in meristem and organ size [33].

The pattern of non-additive gene action was also notable for transcripts associated with cell fate determination, flower development, response to auxin, and seed dormancy (**Fig 2**). These biological functions are well known for having a relevant role in the domestication syndrome [8]. For instance, ZWIP2 is a zinc finger transcription factor that controls organ size by transcriptionally regulating the activity of fruit development-related genes such as *FUL*, *BP*, and *RPL* [34]. *ELF3* is implicated in the regulation of flowering time, a known feature of the domestication syndrome in some crops [35].

Auxin-mediated developmental transitions such as germination and growth are strongly associated with environmental cues. Amongst other genes associated with such functions, there are remarkable examples such as *PKL* and *TOR*. *PKL* is a chromatin remodeling factor whose loss-of-function mutant shows delayed vegetative-to-reproductive transition and overall smaller vegetative and reproductive structures [36].

By repressing the embryonic state and modulating the seed cellular program into a seedling one, *PKL* also determines the transition between the embryonic and vegetative life stage. This process involves cell division, cell expansion, and the activation of photosynthetic machinery [37]. *TOR* is implicated in cell growth and cell division and possibly in seed dormancy [38]. These results highlight the role of non-additive gene action in modulating meristematic activity and ultimately producing the domestication syndrome characterized by gigantism in organs and delayed flowering times.

### Trans-acting factors with high connectivity avoid antagonistic pleiotropy

Natural selection acts on phenotypic variability. Genetic variation in individual genes can produce phenotypic variability. However, cumulative evidence supports an omnigenic model that posits that it is the coordinated action of constellations of genes which ultimately produces phenotypic variation on continuous traits such as organ size [39]. In such constellations or gene regulatory networks (GRNs), the trans-acting elements would produce the majority of the phenotypic variability via expression variation on genes downstream. Therefore, genetic variation at transcription factors and chromatin remodeling factors could be enough to modulate phenotypic variation at the whole GRN level. There is empirical evidence that suggests that to avoid antagonistic pleiotropy, trans-acting factors regulating a large number of genes (large Out-degree) tend to experience small expression changes [40]. Similarly, we reasoned that the number of regulators by which a gene is regulated (In-degree) could produce evolutionary constraint on its expression level, as mutations at some regulators could compensate for the phenotypic effects of other regulators. We hypothesized a negative correlation between the magnitude of expression variation and both, the Out-degree and the In-degree. To formally test this hypothesis, an *A. thaliana* co-expression-based GRN was employed. This GRN represents 330,775 regulatory interactions among 1,883 trans-acting factors (regulators) and 22,478 non-trans-acting genes (targets). From those interactions, 1,828 are self-regulatory. We then obtained the sub-network for a set of 3584 transcripts that display expression differences between the Col and the C24 *A. thaliana* accessions and that had detectable expression in the F1 Col-C24 hybrids. These accessions differ substantially in vegetative growth and reproductive yields [41]. This sub-GRN represents 16,226 regulatory associations between 443 regulators and 3,141 targets, including 416 self-regulatory interactions.

Overall, we found a weak but significant negative correlation between the magnitude of transcript abundance differences (|Log2(Col/C24)|) and both the Out-degree and the In-degree (*ρ* = −0.12, p-val = 0.02; *ρ* = −0.033, p-val = 0.04, respectively) (**Fig 3A left**). These correlation values are similar to those found in a study employing different *Drosophila* species [40].

**Figure 3.**
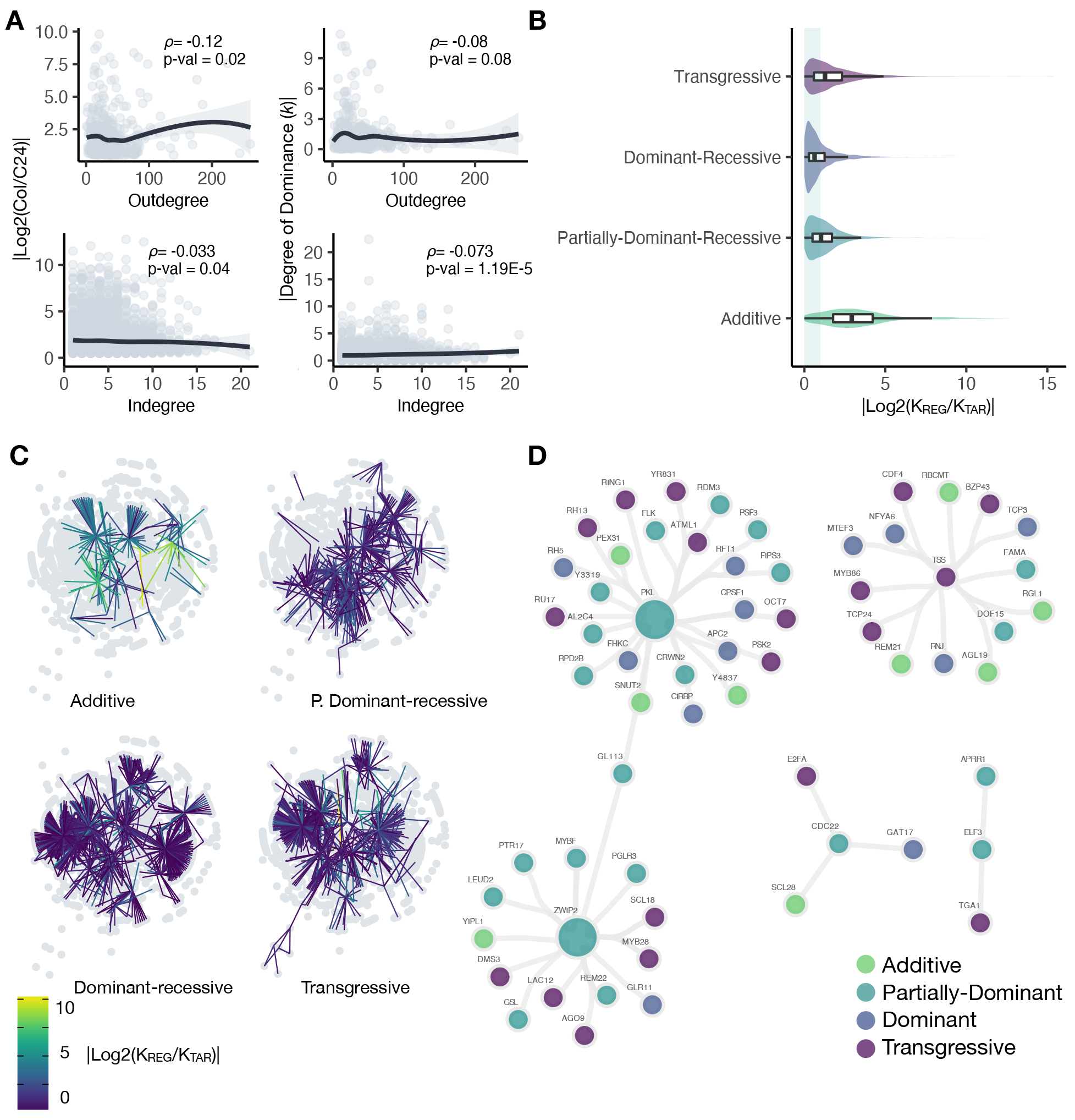
Analysis of the *A. thaliana* GRN showing connectivity properties of regulators and their targets. **A**) The association between the Out-degree and the In-degree to the magnitude of divergence on transcript accumulation between *A. thaliana* Col-0 and C24 as well as their absolute degree of dominance (*k*) are shown as scatter plots. The line depicts the loess regression. **B**) Violin plots showing the absolute Log2 foldchange in the degree of dominance of TFs and their target genes for each category of inheritance. Boxplot depict the median and quartile values. Values of |Log2(K_REG_/K_TAR_)| near to 0 indicate the TF and its target have highly similar values of k. **C**) Gene regulatory network for genes with transcript accumulation divergence between *A. thaliana* Col-0 and C24. Edges are color coded according to the (|Log2(K_REG_/K_TAR_)|). Darker edges denote REG-TAR interaction with high similarity on their degrees of dominance. **D)**Co-expression network for selected regulatory genes. Nodes are color-coded accordingly with the degree of dominance in a discrete fashion.

Recessive phenotypes are often the result of deleterious mutations [42]. So, if we consider the non-additive mode of gene action as a proxy of dominant-recessive variation, it is feasible to predict a negative correlation between the Out-degree and the degree of dominance (*k*). For the case of the In-degree, genes with a large number of regulators could display stronger non-additive effects than their counterparts due to the cumulative effects of their regulators. We tested such hypotheses employing the aforementioned GRN and the degree of dominance estimates in the Col x C24 F1 hybrid. We found a weak but significant negative correlation between the degree of dominance (*k*) and the Out-degree (*ρ* = − 0.08, p-val = 0.08). Conversely, the degree of dominance showed a weak but significant positive correlation with the In-degree (*ρ* = 0.073, p-val = 1.19e-5) (**Fig 3A right**). Despite its relevance, to our knowledge, no previous studies have addressed the association between these two connectivity GRN properties and the degree of dominance.

### Non-additive gene action propagates through GRNs

Under the scenario of GRNs being modulated by the activity of key trans-acting regulators, it is natural to think that different modes of gene action of trans-acting factors would have different phenotypic effects on the genes they regulate downstream. For example, if a trans-acting factor shows non-additive (dominant-recessive) expression in an F1 hybrid background, it would be expected for its target genes to show dominant-recessive gene action. This is because trans-acting factors bind to the two homologous chromosomes in diploid organisms, thus completely recapitulating the phenotype of one parent, unless the trans-acting factor is haploinsufficient. Conversely, for a trans-acting factor showing additive gene action (both functional alleles), it would be expected for its targets to show transgressive gene action because they experience the cumulative trans-acting regulation of the two functional alleles. We tested these hypotheses by establishing a similarity index between the degree of dominance of trans-acting regulators and their targets(|Log2(K_REG_/K_TAR_)|) in our GRN.

In general, we observed that from the total 443 trans-acting regulators contained in the GRN, one third (33.8%) showed transgressive gene action, about one quarter showed dominant-recessive (24.8%) or partially-dominant-recessive gene action (24.8% and 26.6% respectively), and a smaller portion (14.6%) showed additive gene action. This data recapitulates the genome-wide pattern described above for this genetic cross (**Fig 1A**), suggesting that trans-acting variation could be a relevant source of transcriptional variability between the Col and C24 inbred lines. We tested for differences in the similarity index among the four discrete categories of gene action of transcription factors. We found higher similarity in gene action among trans-acting factors and their targets when the trans-acting factor showed dominant-recessive gene action, followed by instances where the trans-acting factor was partially dominant-recessive (median |Log2(K_REG_/K_TAR_)| = 0.63 and 1.03, respectively). These results suggest that trans-acting factors displaying these two modes of gene action tend to produce the same gene action on their targets. Whereas a lower similarity was observed when trans-acting factors showed transgressive gene action (median |Log2(K_REG_/K_TAR_)| =1.28), and even lower for the trans-acting factors showing additive gene action (median |Log2(KTF/K_TAR_)| = 2.93) (**Fig 3B, 3C**). This data indicates, on the one hand, that dominant-recessive allelic interactions are more propagable through GRNs than transgressive gene action. And on the other hand, trans-acting factors showing additive gene action tend to produce transgressive gene action on their targets, supporting our hypothesis of the role of trans-acting variation in heterotic phenotypes. A Wilcoxon rank-sum test in a pair-wise fashion found significant differences in all comparisons. Comparisons that included transcription factors with additive gene action had larger effects sizes (**Table S3**).

The propagation of gene action through GRNs was analyzed in the subnetworks of GRNs of *PKL*, *TSS*, *ZWIP2*, *CDC22*, and *ELF3* (**Fig 3D**). Notably, for *PKL*, which showed partially-dominant-recessive gene action, only three out of its 24 target genes showed additive gene action (binomial test; p = 0.1).

Unfortunately, the size of the other sub-networks is too small to be formally analyzed. These results suggest that genetic variation affecting the activity of trans-acting factors propagates to some degree.

## Discussion

By comparing transcript abundance variation and its patterns of gene action between a single genetic cross of plants evolving under natural selection to those under domestication, we found that the latter exhibit such variation in a substantially larger portion of their transcriptomes and that this variation appears to be driven by LOF genetic mutations that produce non-additive gene action. This result provides evidence of the role of human mediated domestication in shaping gene expression patterns despite an overall small genome-wide genetic divergence compared to species evolving under natural selection. For instance, the switchgrass subspecies exhibit 1.2% of genome-wide nucleotide differences [43], while these values are 0.3%, 0.2% and 0.1% for the chili pepper, maize and sunflower accessions, respectively [44–46].

Despite transcriptional variability due to sampling at varied developmental time points or tissues, our normalized measure of gene action (*k*) allowed us to investigate the genetic nature of allelic variation influencing transcript abundance across different species. Indeed, our study revealed that a substantial portion of the transcripts that showed abundance differences between wild and domesticated plants display partially non-additive (dominant-recessive) gene action. This mode of gene action has been ignored in the majority of experiments assessing the inheritance of transcript accumulation in plants [19–21, 24, 29, 47–50]. Perhaps due to the use of discrete categories of modes of gene action based on differential expression between F0s and F1 hybrids. Our findings suggest a relevant role of non-additive gene action on modulating a polygenic architecture of domestication-related phenotypes.

The recessive patterns of transcript accumulation observed in the domesticated sunflower and chili pepper suggest that divergence due to selection under domestication is associated with loss-of-function genetic variation. The opposite pattern was shown in maize crosses, which indicates a possible role of gain-of-function genetic changes in the domestication of this crop. However, it is likely that maize-like alleles initially arose in a teosinte genetic background and behaved as loss-of-function. So the dominance of modern maize phenotypes that we observe in an F1 hybrid could differ from the initial conditions of maize domestication [51]. This finding supports a growing collection of studies that propose human-mediated selection for gigantism of organs, unconsciously selected for genetic variation that would not be selected in naturally-evolving populations due to its deleterious effects on fitness [42].

The gigantism of vegetative and reproductive organs is a common feature of domesticated plants. Several studies have proposed a relevant role of loss-of-function variation in cell cycle and cell fate determination in producing such gigantism by altering meristem size [2, 8]. Congruently, our study revealed that transcript abundance of genes associated with cell cycle and cell fate determination showed dominant-recessive gene action. Empirical evidence indicates that genes involved in such functions belong to homologous GRNs, suggesting crop domestication is an outstanding example of parallel evolution. For instance, the modification of plant architecture and inflorescence via the modulation of the cell cycle GRN by the action of the TCP transcriptional repressor *tb1* is common for maize, pearl millet, and barley [3]. In our study, both *tb1* and transcripts annotated as cell cycle regulators showed strong non-additive effects in the crosses of maize, chili pepper, and sunflower. We could hypothesize that if GRNs are at least partially conserved between these species, the non-additive trans-regulatory activity of their *tb1* homologs could contribute to alterations in the cell cycle machinery and meristem size. It would be interesting to experimentally test the aforementioned hypothesis by contrasting the meristem size of both gain and loss-of-function mutants of *tb1* in different species.

Using an *A. thaliana* GRN and differential expression data of two accessions of this species that differ in meristem size, we found a weak negative correlation between the magnitude of expression divergence and both, the In-degree and the Out-degree. These findings suggest that trans-acting factors with high Out-degree avoid antagonistic pleiotropy by constraining their expression levels. Whereas the constraint of expression differences on genes with high In-degree could be due to the compensatory regulation of their multiple trans-acting regulators [40]. Our observation of a weak negative correlation between the degree of dominance (*k*) and the Out-degree has no previous record in the literature. We speculate, however, that given that mutations producing non-additive phenotypes often result in negative effects on fitness [42], natural selection would purge such deleterious mutations on highly connected trans-acting regulators to avoid antagonistic pleiotropy. It remains to be seen if our findings are supported by GRNs of other domesticated species, yet, they add to our knowledge about the interplay between GRNs and the nature of genetic changes controlling domestication-related phenotypes such as organ size. Finally, we acknowledge the use of *Arabidopsis* microarray-based expression profiles to build the co-expression based GRN could be a relevant source of technical bias. Yet this data set is a valuable resource for the plant biology community.

The nature of genetic changes that produce non-additive gene action within GRNs at two levels (dominant-recessive and transgressive), led us to consider two classical hypotheses that propose mechanistic explanations [17, 52]: *i*) the dominance hypothesis posits that the deleterious allelic effects of one parent are complemented in the F1 hybrid by the functional allele of the other parent. *ii*) The beneficial allele hypothesis predicts that two functional alleles would produce a transgressive phenotype in the F1 hybrid due to their complementary beneficial phenotypic effects. Our study provides relevant empirical evidence for these two hypotheses. First, in support of the dominance hypothesis, we found that trans-acting factors showing dominant-recessive gene action produce dominant-recessive effects on most of their target genes. In agreement with this finding, computational models propose that dominant-recessive gene action propagates through GRNs via trans-acting factors [18]. One possible explanation for this finding is that, in absence of cis-acting variation in the target genes, the functional trans-acting allele complements the non-functional trans-acting allele of the other parent. Secondly, in support of the beneficial allele hypothesis, our data revealed that targets genes of trans-acting factors that showed additive gene action tended to display transgressive phenotypes. This indicates that parental allelic trans-acting factors differ in their cis-acting regulatory elements, but the two alleles are fully functional in their respective genetic background, thus reinforcing their regulatory activity on their transcriptional targets.

The beneficial allele hypothesis is an example of transcriptional epistasis. A classic example of transcriptional epistasis is the anthocyanin pathway in maize, where the additive interaction of allelic variants of the trans-acting regulator *B*, produces transgressive transcript abundance on the anthocyanin synthesis genes *A1*, *A2*, and *Bz1* [53]. The catalog of 65 trans-acting regulators that showed additive inheritance in the Col-C24 *A. thaliana* cross is a relevant source of candidates to experimentally test the role of functional cis-acting allelic variants in producing transgressive gene action on meristem size. It remains to be seen if our findings are supported by GRNs of other species, yet, they add to our knowledge about the interplay between GRNs and the nature of genetic changes controlling domestication-related phenotypes such as organ size.

Finally, our finding in *A. thaliana* of the propagation of phenotypic effects of trans-acting regulators to their targets adds empirical evidence to the omnigenic model. According to this model, variability in complex phenotypes such as organ size is the result of variations on the trans-acting activity of “peripheral” TFs on the expression of “core” genes in GRNs. Therefore, the domestication-related phenotypes are likely underlaid by a small number of mutations that work in an omnigenic fashion [39]. As large transcriptomic datasets are available to construct high-quality GRNs it will be possible to test our hypothesis in other plant species.

## Conclusions

Our study provides evidence of the role of non-additive gene action in the modulation of transcript abundance of genes associated with domestication-related traits in plants, such as the gigantism of reproductive organs. The non-additive transcript abundance variation on genes associated with cell cycle and cell fate determination machinery are likely driven by loss-of-function mutations, in line with the classic observation of aberrant organ sizes in domesticated plants made by Darwin. One major limitation of this study is the use of a single genetic cross as natural selection control. It remains to be seen if the use of additional genetic crosses as tests and as controls, would shed light on whether non-additive patterns of gene expression play a major role in plant domestication. Using a co-expression-based GRN of the model plant *A. thaliana*, our data revealed a weak but significant negative correlation between the Out-degree and both the magnitude of transcript abundance variation a well the absolute degree of dominance. Which suggested highly connected genes in GRNs are less prone to be influenced by LOF genetic changes that result in non-additive gene action. If the Out-degree is a good measure of pleiotropy, LOF mutations would be expected to be preferentially eliminated by natural selection, providing a mechanism to avoid antagonistic pleiotropy. Finally, our study provides empirical evidence of the propagation of non-additive phenotypes in GRNs. This is congruent with the omnigenic model and highlights the role of trans-acting regulatory changes in modulating polygenic phenotypes such as fruit size in the evolution of plants under domestication.

## Methods

### Analysis design, data collection and preprocessing

We reasoned that F1 hybrids derived from crosses between cultivated plants and their wild progenitors would display a large portion of non-additive variation in their transcriptional landscapes compared to genetic crosses between lineages evolving under natural selection. Therefore, our dataset includes one genetic cross of two switchgrass (*Panicum hallii hallii* x *P. hallii filipes*) ecotypes evolving under natural selection [29]; three genetic crosses of cultivated maize and its wild progenitors (*Zea mays* x *Zea mays parviglumis*) [22]; one genetic cross of cultivated and wild chili pepper (*Capsicum annuum annuum* x *C. annuum glabriusculum*). [9]; one genetic cross of cultivated and wild sunflower (*Heliantus annuus* x *H. petiolaris*) [19]; one reciprocal cross between two inbred lines of cultivated rice (*Oriza sativa indica* x *O. sativa japonica*) [54]; and two reciprocal crosses of the *Arabidopsis thaliana* Columbia to the *A. thaliana* C24 and *A. thaliana* Landsberg *erecta* ecotypes [55], which are domesticated to the laboratory conditions. Due to the availability of raw next generation sequencing NGS-based transcriptome data, reference transcriptomes and the experimental conditions from which RNA was obtained, data for specific genetic crosses was chosen and processed into a unified analytical custom pipeline (**Table S4**). Raw NGS sequences were retrieved and converted to fastq format employing the fasterq-dump program of the SRA toolkit 2.9.1 [56] using default parameters. Reads were then pre-processed to remove adapters, and to trim and filter low quality bases using the fastp software [57] with default parameters.

### Read mapping, transcript accumulation quantification and differential expression analyses

Genome-wide transcript abundance quantification for each triplet (Both parents + F1 hybrid) of sequencing libraries was estimated by pseudo-aligning the clean reads to their respective reference transcriptome using Kallisto [58] with parameters −b 100, −t 16 and -bias. Data for each species was then consolidated into a single data base that included both parents (F0) and their (F1) hybrid. Transcripts with a per-species median of read counts smaller than five were excluded for further analyses. Transcript abundance estimates were then employed to test for differential transcript abundance between F0 genotypes using a generalized linear model via the DESeq2 R library [59]. Transcripts were considered as differentially expressed using an FDR ≤ 0.05 and a |Log2 fold change| > 0.5.

### Mode of gene action assessment

We estimated the degree of dominance (*k*) as a proxy of mode of gene action for all the transcripts showing evidence of both differential expression between F0 and identifiable expression in the F1 hybrids. Specifically, the degree of dominance (*k* = d/a) estimates the association between the dominant and additive effects. As we see it, the degree of dominance is a normalized continuous measurement that estimates the deviation of the null hypothesis of both parents contributing equally to the observed phenotype in the F1 hybrids. The dominance effect (d = F1 − [(X_1_ + X_2_)/2]) measures the deviation of the F1 hybrid phenotype from a midpoint of the F0s. The additive effect (a = (X_1_−X_2_)/2) provides a quantitative estimate of the effect of substituting one allele, for example a wild allele for a mutant allele in a cultivated phenotype. Using a custom R function, we employed the normalized transcript abundance values and evaluated *k* for all the differentially expressed transcripts across 12 genetic crosses. We then sorted each transcript according to four discrete categories of gene action as follows: |k| < 0.25 = additive; |*k*| ≥ 0.25 & |*k*| ≤ 0.75 = partially dominant-recessive; |*k*| ≥ 0.75 & |*k*| ≤ 1.25 = dominant-recessive; |*k*| ≥ 1.25 = transgressive. Beyond discrete categories of transcript abundance gene action, we analyzed the transcriptional landscapes employing probability distributions. This allowed us to compare the breath of allelic interactions in different contexts of selective pressures.

Genetic crosses were then split into Natural Selection, Domestication and Inbreds according to the publications from where data was retrieved (supplementary table 1). We then performed a Kolmogorov-Smirnov test in a pair-wise fashion to asses if plants evolving under natural selection accumulate more additive variation than plants evolving under domestication. Data were then visualized using an empirical cumulative distribution function.

### Gene Ontology and Co-expression network construction

To obtain biological insight into the portion of the transcriptome showing differential expression and showing different modes of gene action, we employed GO enrichment and co-expression network analyses. To avoid possible annotation artefacts, for instance ambiguity in the protein domains and biological functions, we decided to generate a unified annotation for the set whole set of differentially expressed transcripts. We first blasted [60] the transcripts against the Swiss-Prot database [61] and then selected the best hit employing home-made R scripts. GO terms for each transcript were then retrieved from the UniProt database using the ID mapping function [62]. The rationale of our GO enrichment analysis was to test if genes in different categories of gene action in each cross are enriched for different biological functions. The topGO R package [63] was then used to perform the enrichment analysis employing the weight01 algorithm.

Gene co-expression networks allow for inferring groups of genes that act together to regulate cellular processes. While some widely used approaches such as the WGCNA [64] employ correlation as a gold standard measurement of gene co-expression, other algorithms such as AracNe-AP [65] have been shown to perform better to infer regulatory relationships such as those between transcription factor genes and non-transcription factor genes. Using the AracNe-AP algorithm and 79 public microarray-based transcriptomes across different organs and developmental stages [66], we thus inferred a co-expression-based gene regulatory network (GRN) for the model organism *Arabidopsis thaliana*. Regulatory associations between TFs and non-TF genes were computed using the mutual information measurement. The AracNe-AP algorithm takes a list of genes defined as TFs and the rest of the genes as possible targets and infers mutual information. TFs are connected by an edge in a network if the mutual information meets a threshold value estimated from random sampling of mutual information among TFs and non-TF genes. We then obtained the specific co-expression or regulatory network for the genes showing differential gene expression between the *A. thaliana* Col-C24 F0s.

### Association between Gene Regulatory Network properties, transcript abundance differences and gene action

Some TFs have thousands of transcriptional targets downstream, it has been reported that there is a negative correlation between the effect size of transcriptional variation (ES) at TFs and the number of genes they regulate (Out-degree). In a similar way, the number of TFs that regulate a single gene (In-degree) has been negatively be correlated with the ES. We thus employed a Pearson correlation analysis and a Loess non-linear regression to investigate the possible association between ES and both the Outdegree and the Indegree.

We were also interested in assessing the association between both Indegree and Outdegree to the degree of dominance (*k*). We predicted that TFs with high Outdegree would tend to show additive gene action (|*k*| ~ 0). We have no clear prediction for the inheritance of genes with high Indegree. We employed correlation and non-linear regression analyses to investigate the association between those network metrics and the degree of dominance (*k*).

### Gene action propagation in Gene Regulatory Networks

We reasoned that TFs and their targets would display similar degrees of dominance (*k*), particularly for TFs showing partial or complete dominant-recessive gene action. We thus employed the absolute values of the Log2 fold change between the *k* values of TFs and their targets (|Log2(K_REG_/K_TAR_)|), to generate a normalized measurement of degree of dominance similarity. In addition to visualize the distributions of k similarity among the four categories of gene action of TFs, we explicitly compared the median similarity of TFs showing additive gene action to those showing non-additive gene action using a Wilcoxon rank sum test using the wilcox_test of the Rstatix R package [67]. We then mapped the k similarity values as a color-coded edge attribute and visualized the co-expression networks for each instances of gene action of TFs using the ggraph R package [68]. To visually analyze the similarity of degree of dominance between TFs and their targets, networks for a set of TFs annotated as cell cycle, cell fate and response to auxin regulators were plotted. All data visualizations and data wrangling, unless specified, were done using the Tidyverse R package [69] in the Rstudio programming environment [70, 71].

## List of abbreviations

APC: Anaphase-promoting complex
CDC22: Cell division cycle 20.2
CDKD1: cyclin-dependent kinase D1
DE: Differentially expressed
ELF3: Early flowering protein
ES: Effect size
FC: Fold change
FDR: False discovery rate
GO: Gene ontology
GRN: Gene regulatory network
NGS: Next generation sequencing
PKL: Pickle remodeling factor
TOR: Target of rapamycin
ZWIP2: Zink finger protein WIP2

## Declarations

### Ethics approval and consent to participate

Not applicable

### Consent for publication

Not applicable

### Availability of data and materials

The datasets generated and analyzed during the current study are available from the figshare repository: https://doi.org/10.6084/m9.figshare.19358249.v1

### Authors’ contributions

E.D-V. and A.C-J. conceived and designed the project. E.D-V and D.H-R. performed the computational analyses. E.D-V. wrote the article. A.C-J. critically revised the article.

## Acknowledgements

E.D-V. is grateful to Dulce Valdivia for her help to improve some R scripts and for her time to discuss the statistical approaches.

## Supplementary Figures

**Figure S2.**
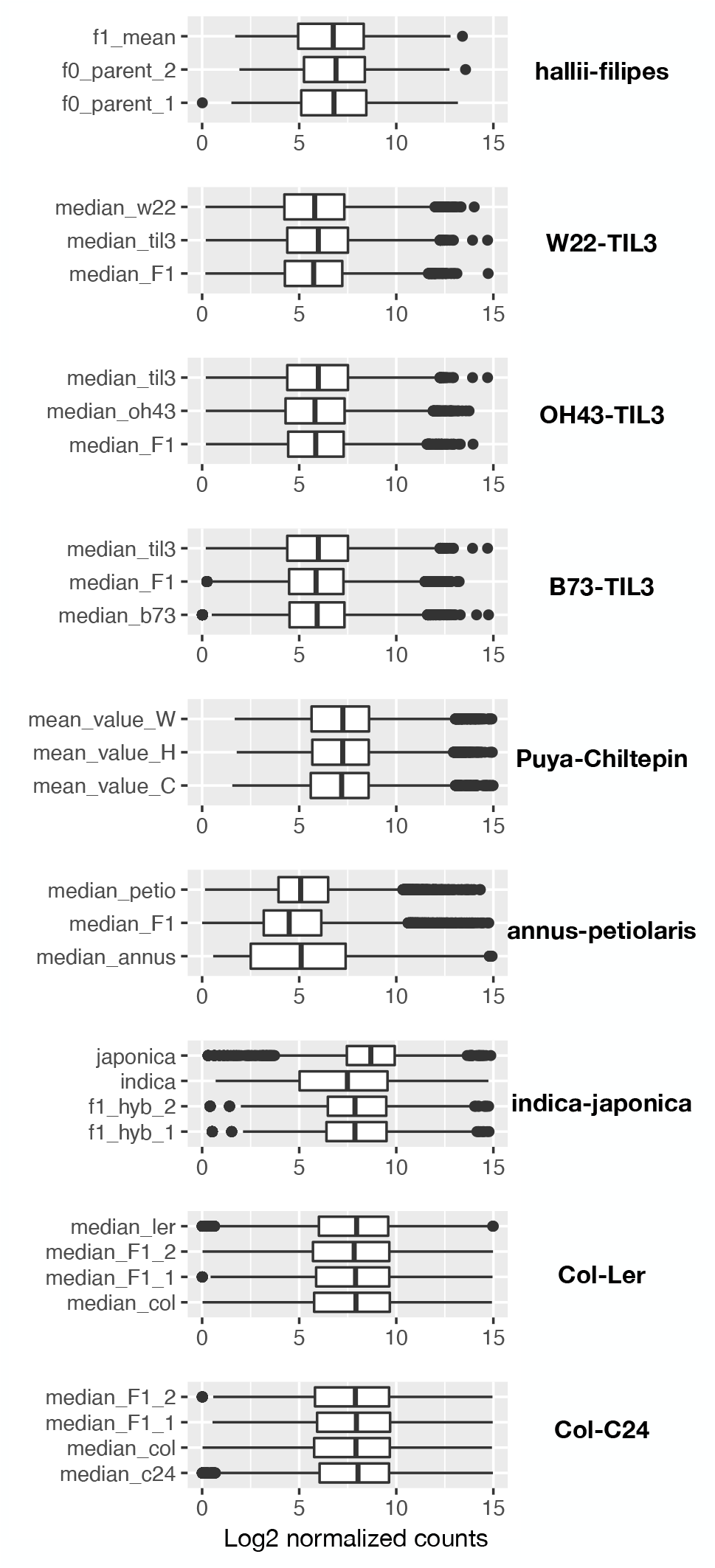
A sampling-based analysis revealed a uniform ~60X sequencing coverage for all the libraries used in the study. The per-transcript sequencing coverage (Log2 normalized read counts) is shown as boxplots for the three genotypes of each species. The line within the boxes depicts the median, whereas the dots outside the box represent outlier values.

**Figure S1.**
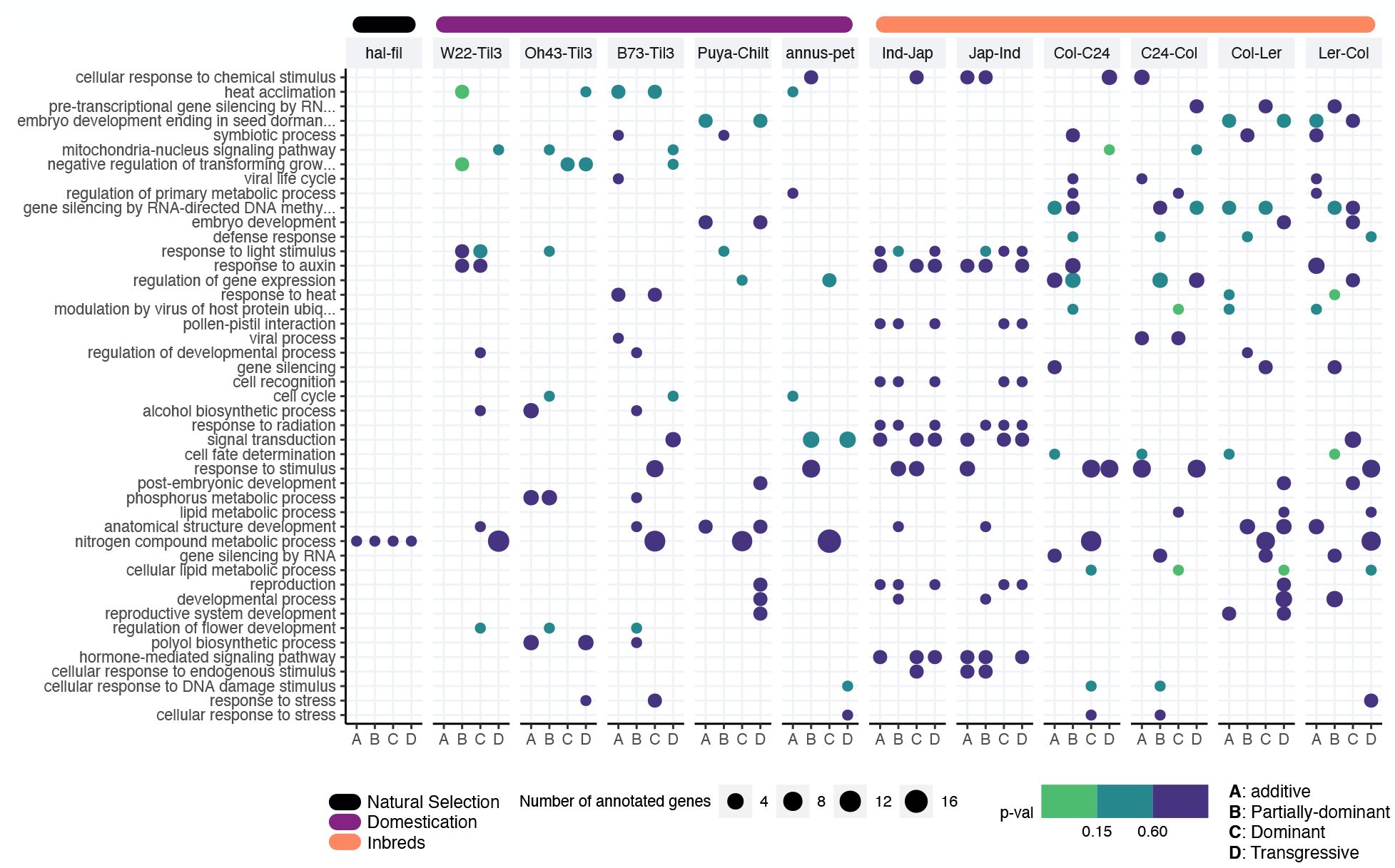
GO enrichment analysis showing biological functions altered in the studied genetic crosses. Dot plots show the enrichment, or presence of each GO for each mode of gene action (Additive, Partially-Dominant, Dominant, Transgressive) across the array of genetic crosses. Color of dots indicate the significance of the enrichment; the size depicts the number of annotated genes to each GO.

**Table S1.**
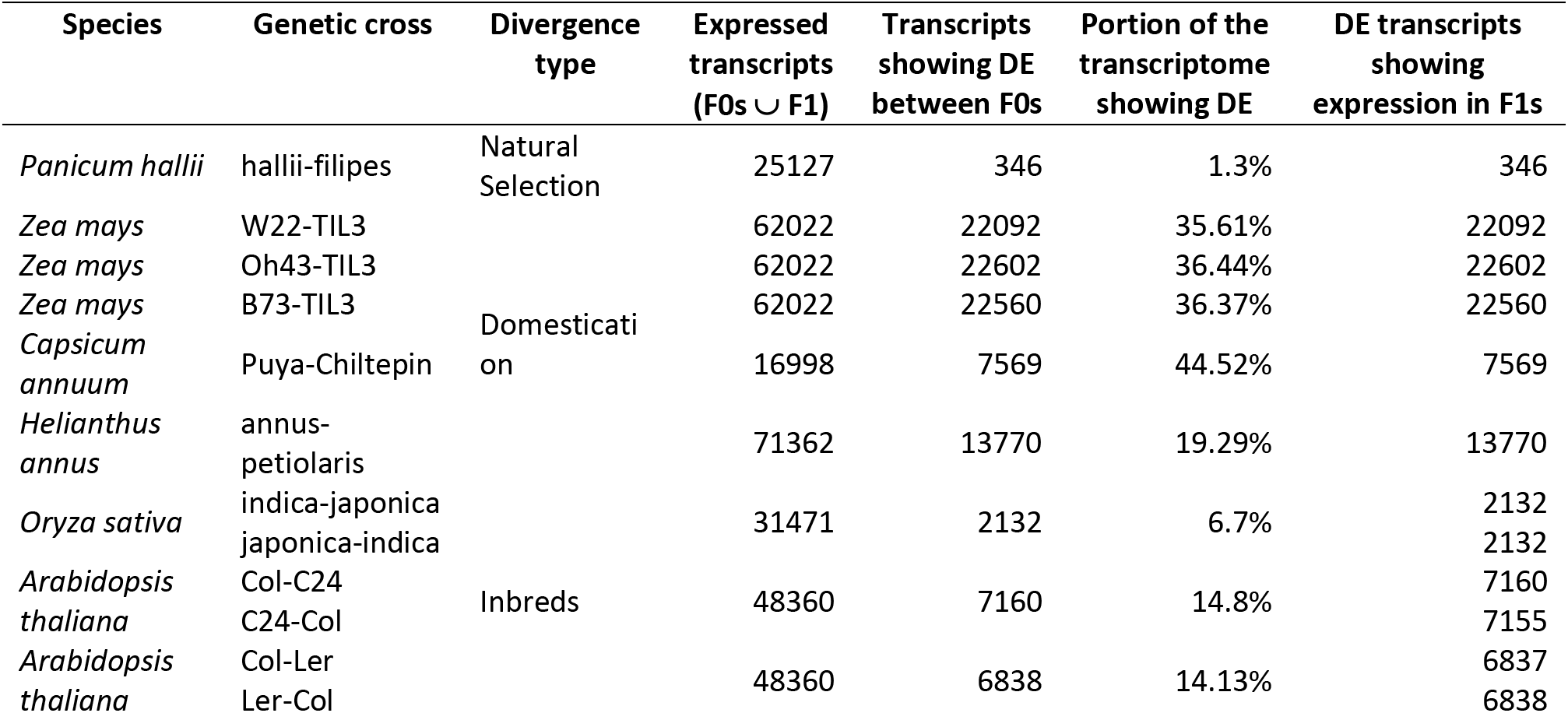

**Table S2.**

| Biological function | Divergence type | Number of transcripts | Median $\vert k\vert $ | p-val Wilcoxon $\mu \vert k\vert > 0.25$ |
| --- | --- | --- | --- | --- |
| Cell cycle | Natural Selection | 1 | NA | NA |
|  | Domestication | 171 | 0.8 | 3.73e-113 |
|  | Inbreds | 834 | 0.76 | 1.20e-26 |
| Cell fate | Natural Selection | 0 | NA | NA |
|  | Domestication | 22 | 0.41 | 0.0264 |
|  | Inbreds | 14 | 0.56 | 0.0262 |
| Flower development | Natural Selection | 4 | NA | NA |
|  | Domestication | 350 | 0.91 | 1.022-48 |
|  | Inbreds | 121 | 0.79 | 3.40e-20 |
| Response to auxin | Natural Selection | 3 | NA | NA |
|  | Domestication | 645 | 0.9 | 2.79e-91 |
|  | Inbreds | 239 | 0.78 | 2.30e-33 |
| Seed dormancy | Natural Selection | 3 | NA | NA |
|  | Domestication | 1067 | 0.78 | 4.36e-135 |
|  | Inbreds | 254 | 0.76 | 1.23e-37 |

**Table S3.**

| TFs group 1 | TFs group 2 | TFs group 1 | TFs group 2 | Interactions group 1 | Interactions group 2 | Effect size | Adjusted p value |
| --- | --- | --- | --- | --- | --- | --- | --- |
| Additive | Dom-Rec | 65 | 110 | 2107 | 4555 | 0.58 | ~ 0 |
|  | P. Dom-Rec |  | 118 |  | 4513 | 0.51 | ~ 0 |
|  | Transgressive |  | 150 |  | 5051 | 0.39 | 4.44e-238 |
| Dom-Rec | P. Dom-Rec | 110 | 118 | 4555 | 4513 | 0.19 | 4.44e-77 |
|  | Transgressive |  | 150 |  | 5051 | 0.29 | 2.29e-179 |
| P. Dom-Rec | Transgressive | 118 | 150 | 4513 | 5051 | 0.12 | 1.61e-35 |

**Table S4.**
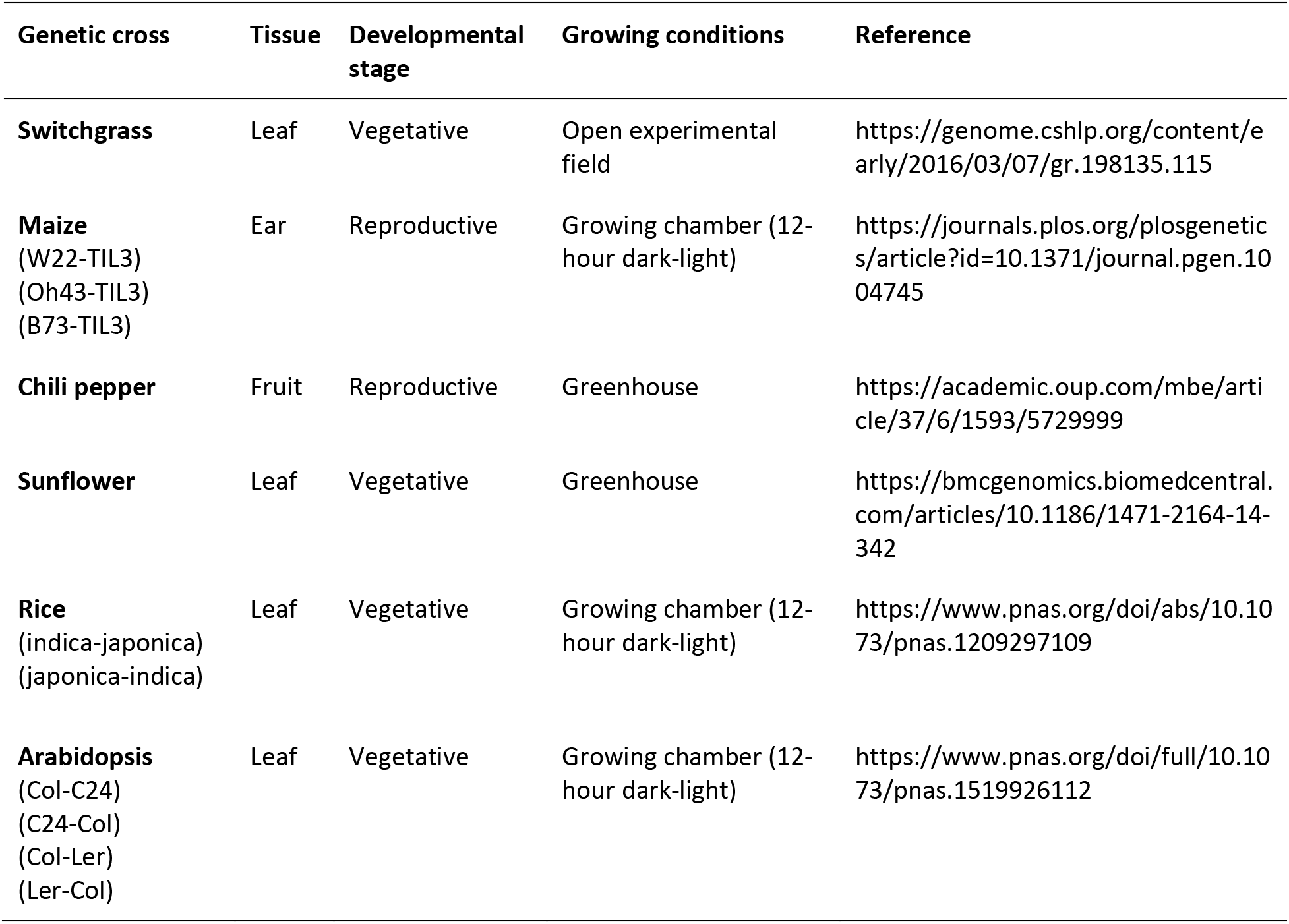

